# Resolving the insertion sites of polymorphic duplications reveals a *HERC2* haplotype under selection

**DOI:** 10.1101/488726

**Authors:** M. Saitou, O. Gokcumen

## Abstract

Polymorphic duplications in humans have been shown to contribute to phenotypic diversity. However, the evolutionary forces that maintain variable duplications across the human genome are largely unexplored. To understand the haplotypic architecture of the derived duplications, we developed a linkage-disequilibrium based method to detect insertion sites of polymorphic duplications not represented in reference genomes. This method also allows resolution of haplotypes harboring the duplications. Using this approach, we conducted genome-wide analyses and identified the insertion sites of 22 common polymorphic duplications. We found that the majority of these duplications are intrachromosomal and only one of them is an interchromosomal insertion. Further characterization of these duplications revealed significant associations to blood and skin phenotypes. Based on population genetics analyses, we found that the partial duplication of a well-characterized pigmentation-related gene, *HERC2*, may be selected against in European populations. We further demonstrated that the haplotype harboring the partial duplication significantly affects the expression of the *HERC2P9* gene in multiple tissues. Our study sheds light onto the evolutionary impact of understudied polymorphic duplications in human populations and presents methodological insights for future studies.

## INTRODUCTION

Genomic copy number variation (duplications and deletions of sections of the genome) has increasingly been appreciated as a driver of human phenotypic variation, accounting for several key adaptive phenotypes, such as HIV resistance (Sabeti et al. 2005), malaria resistance (Leffler et al. 2017), among others. In parallel, several studies have found new connections between copy number variants and disease susceptibility (Zhang et al. 2009; Weischenfeldt et al. 2013). One of the best-known examples of the evolutionary impact of copy number variation on phenotype is the dramatic expansion of *AMY1* copy number in the human lineage, likely driven by high starch consumption (Meisler & Ting 1993). In fact, the gene duplications of *AMY1* remain polymorphic in humans, correlating with historical starch consumption among different human populations (Perry et al. 2007). Despite their genomic and phenotypic impacts, few studies have addressed the evolutionary forces that shape the evolutionary trajectories of polymorphic duplications. We argue that the main reason for the paucity of evolutionary studies on polymorphic duplications is that current, short-read based discovery and genotyping approaches are unable to resolve the genomic locations of inserted duplicated gene copies; consequently, the haplotypic variation associated with a given duplication cannot be fully studied. This problem is further aggravated by the complexity of a considerable portion of the loci harboring duplications, *i.e*., they involve highly repetitive sequences (Sudmant et al. 2015).

Here, by applying a novel linkage disequilibrium based method to the 1000 Genomes Phase 3 Dataset (Sudmant et al. 2015), we detected the insertion sites of 22 common human polymorphic duplications. This dataset allowed us to more thoroughly investigate the potential adaptive contributions of some of these duplications on human phenotypic diversity.

## RESULTS

### Detecting putatively adaptive duplications and their insertion sites

Short-read sequencing technologies can use read-depth and, for certain occasions, paired-end analysis to detect polymorphic duplications as compared to reference genomes (Mills et al. 2011). However, these methods are limited in their ability to detect the insertion sites and hence the haplotypes harboring these duplications. This information is crucial to conduct neutrality tests and functional analyses.

To detect insertion sites of polymorphic duplications, we utilized genome-wide linkage disequilibrium between the genotyped duplications (for which the insertion site is unknown) and single nucleotide variations across the human genome. Specifically, we assumed that when a duplicated sequence was inserted in a certain genomic region and subsequently increased in allele frequency, the flanking single nucleotide variants would show linkage disequilibrium with the duplication (**fig. 1A**). This method can only detect the insertion sites of gene duplications with relatively high allele frequency. The signal weakens considerably if the haplotype harboring the duplicated sequence undergoes recombination or gene conversion. It is also important to note here that our method’s power depends on the accuracy of variation calls and phasing of the database that we are using.

**FIG. 1.**
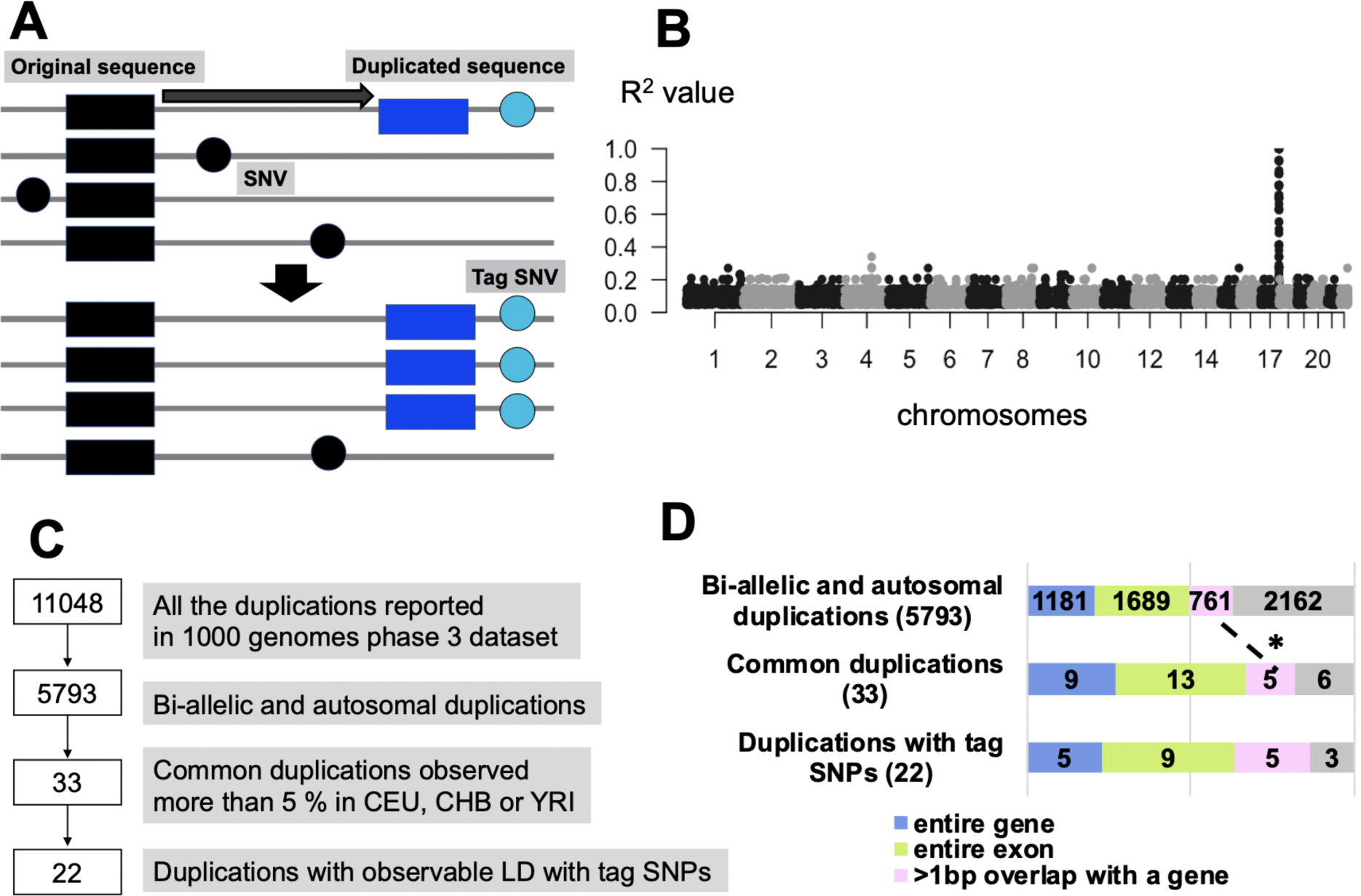
The strategy of the linkage disequilibrium-based method to detect the insertion region of the polymorphic duplication. **(A)** The schematic representation of our approach to detect insertion sites and haplotypes harboring polymorphic duplications. **(B)** One example of the linkage disequilibrium-peak between and duplication and single nucleotide variants. Each dot indicates a single nucleotide variant. The X-axis shows chromosomal locations and Y-axis shows the linkage disequilibrium between each single nucleotide variant and a specific polymorphic duplication, in this case, esv3641421, in the CEU population. **(C)** The filtering process of the duplications. **(D)** The breakdown of the number of duplications based on their exonic content and allele frequency. The legend below indicate the color-coded functional categories. We observed the enrichment of genic duplication in the common duplications compared to the initial duplication set (p-value = 0.03684, one-tail Pearson’s Chi-squared test with Yates’ continuity correction)

We chose to apply this method to the data provided by the 1000 Genome Project phase 3 dataset (Sudmant et al. 2015), which includes 11,048 variable human duplications. We chose this dataset as it remains the most accurate population-level compilation of human variable duplications and phased single nucleotide variants essential for our analysis. To minimize false-positive calls and to avoid complicating our dataset, we conducted preliminary filtering (**fig. 1B**). First, we eliminated multiallelic copy number variations and focused only on bi-allelic duplications reported as 2, 3, or 4 diploid copies in humans. To increase our power for detecting linkage disequilibrium, we focused on common duplications observed in more than 5% in any of Central Europeans from Utah (CEU), Yoruba from Ibadan (YRI), or Han Chinese from Beijing (CHB). After this filtering, we were left with 33 common, bi-allelic duplications for this study. We manually identified observable peak(s) of linkage disequilibrium for 22 out of 33 common duplications (**fig. 1C**) with single nucleotide variations across the genome (**Table 1, Table S1**).

We found that 21 out of 22 (~ 96%) of duplication insertion sites are found on the same chromosome as where the duplicated sequence is found. Further scrutinization of the haplotypes harboring intrachromosomal duplications revealed that five of them overlap with the original duplicated gene, six of them were located (>1kb) upstream of the region and eight of them located (> 1kb) downstream of the region (**Table 1 and fig. S1**). Additionally, we found that one duplication (esv3631000) containing the gene *ZNF664*, inserted into chromosome 2. This observation was supported by 17 SNPs on chromosome 2 in strong linkage disequilibrium (R^2^ > 0.8) with the duplication, which is originally on chromosome 12 (**Table S1**). *ZNF664* is classified as retro-duplication (Abyzov et al. 2013). Thus, the retroposon machinery may facilitate a copy and paste mechanism of the reverse transcribed mRNA of the original gene to a random insertion point, in this case, chromosome 2.

### The functional impact of haplotypes harboring common duplications

We then scrutinized the genic content of the filtered duplications. Of the 22 duplicated sequences, 5 contain whole genes, 9 contain coding exonic sequences, 4 contain intronic sequences, and 3 contain only intergenic sequences (**Table 1**). We then asked whether the high proportion of duplications containing coding sequences is more than expected, especially given that a previous study reported that only ~20% of common duplications overlap with coding sequences (Conrad et al. 2010). This contrasts with the more than 50% of duplications we outlined that overlap with an entire gene or coding exon (**fig. 1D**). We found that duplications associated with strong linkage disequilibrium single nucleotide variants do not significantly differ in their coding sequence content from duplications that do not (**fig. 1D**). Surprisingly, we further found that 1000 Genomes Phase 3 duplications show an increase in exonic content with higher allele frequency, contrasting with prior studies where common duplication alleles tended to harbor less coding sequence content than rare duplication alleles (Conrad et al. 2010). This difference may be due to variation in the accuracy and sensitivity of duplication discovery methods, and further underline the necessity for a better scrutinization of copy number variation in human genomes.

Our approach resolved the haplotype architecture of additional gene duplications that may have important functional consequences (**Table 1**). The ascertainment bias in most functional databases limits further scrutinization of the functional impact of polymorphisms to some extent. Specifically, most comprehensive datasets for expression quantitative trait loci analysis and most genome-wide association studies (e.g., UK Biobank) were constructed mostly by data gathered from western European individuals. Majority of gene duplications for which we were able to resolve the haplotypes for were found in African populations only (**Table 1**). Still, we were able to search for specific associations of 8 gene duplication haplotypes with > 5% allele frequency in European populations with gene expression levels documented in GTEx (Lonsdale et al. 2013), as well as with 77 traits documented in UK Biobank (Canela-Xandri et al. 2018). We found 2 significant associations. First, we found the exonic duplication involving *TEX19* gene (esv3645658, tag-variant: rs74001624 (R^2^=1)) is significantly associated with lower levels of expression of the *SECTM1*. Further, we conducted a similar search in the UK-Biobank Phenome-Wide Association Study database and found that the duplication haplotype is significantly (*p*=2.828×10^−16^) associated with Monocyte percentage (**Table 1**). This finding is concordant with the previous findings that *SECTM1* is involved in hematopoietic processes (Slentz-Kesler et al. 1998).

Second, we found that the haplotype harboring esv3585141 duplication, which involves non-coding sequences only (tag-variant: rs74865018 (R^2^=0.5), is associated with skin color in UK-Biobank Phenome-Wide Association Study database (*p*=1.373×10^−9^). However, we have not found a significant association with the expression levels of any neighboring genes based on our search in the GTex database. Our previous research has shown that copy number variants, including gene duplications, may be important factors in shaping skin/hair phenotypes (Eaaswarkhanth et al. 2014, 2016; Pajic et al. 2016). Indeed, we found that one of the haplotypes in our study harbors the whole gene duplication of the *KRT34* (RefSeq: NM_021013), a member of the keratin gene family, which is important for hair phenotypes and is shown to be affected by copy number variation. This whole gene duplication is common in African populations, but not observed in Eurasian populations (**Table 1**). Gene expression of *KRT34* in human hair follicles is higher in young individuals than that in old individuals (Giesen et al. 2011). Based on GTEx data (Lonsdale et al. 2013), we demonstrate that the haplotype harboring the duplication led to an increase in the dosage of the *KRT34* expression. Interestingly, this duplication shows high linkage disequilibrium (R^2^=0.80) with the adjacent deletion of the *KRT33B* (esv3640584, chr17:39506753-39525903), which may suggest that *KRT34* replaced *KRT33B* through gene conversion.

Overall, resolving the haplotypes harboring gene duplications provide a powerful framework to further scrutinize the functional impact, if any, of these variants. Our observations involving the highlighted genes provides candidates for future evolutionary and biomedical studies.

### Partial *HERC2* duplication may be selected against in European populations

Our main goal in this paper is to leverage the haplotypes of the polymorphic duplications to identify potential selective forces acting on specific polymorphic duplications. To achieve this, we first calculated the allele frequency differences between populations for the 22 polymorphic duplications that we focus in this study. Then we compared these differences to those calculated for randomly selected 3,102 common (>5% alternative allele frequency in CEU, YRI or CHB to match our initial filtering) single nucleotide variants extracted from 1000 Genomes Phase 3 dataset (The 1000 Genomes Project Consortium et al. 2015) (**fig. 2A**). We found that a partial duplication of a well-characterized gene, the HECT And RLD Domain Containing E3 Ubiquitin Protein Ligase 2 gene (*HERC2*) showed apparently higher allele frequency differentiation from the other gene duplications as well as the majority of random single nucleotide variants analyzed as a null background (**Table 1, fig. 2AB, fig. S2**).

**FIG. 2.**
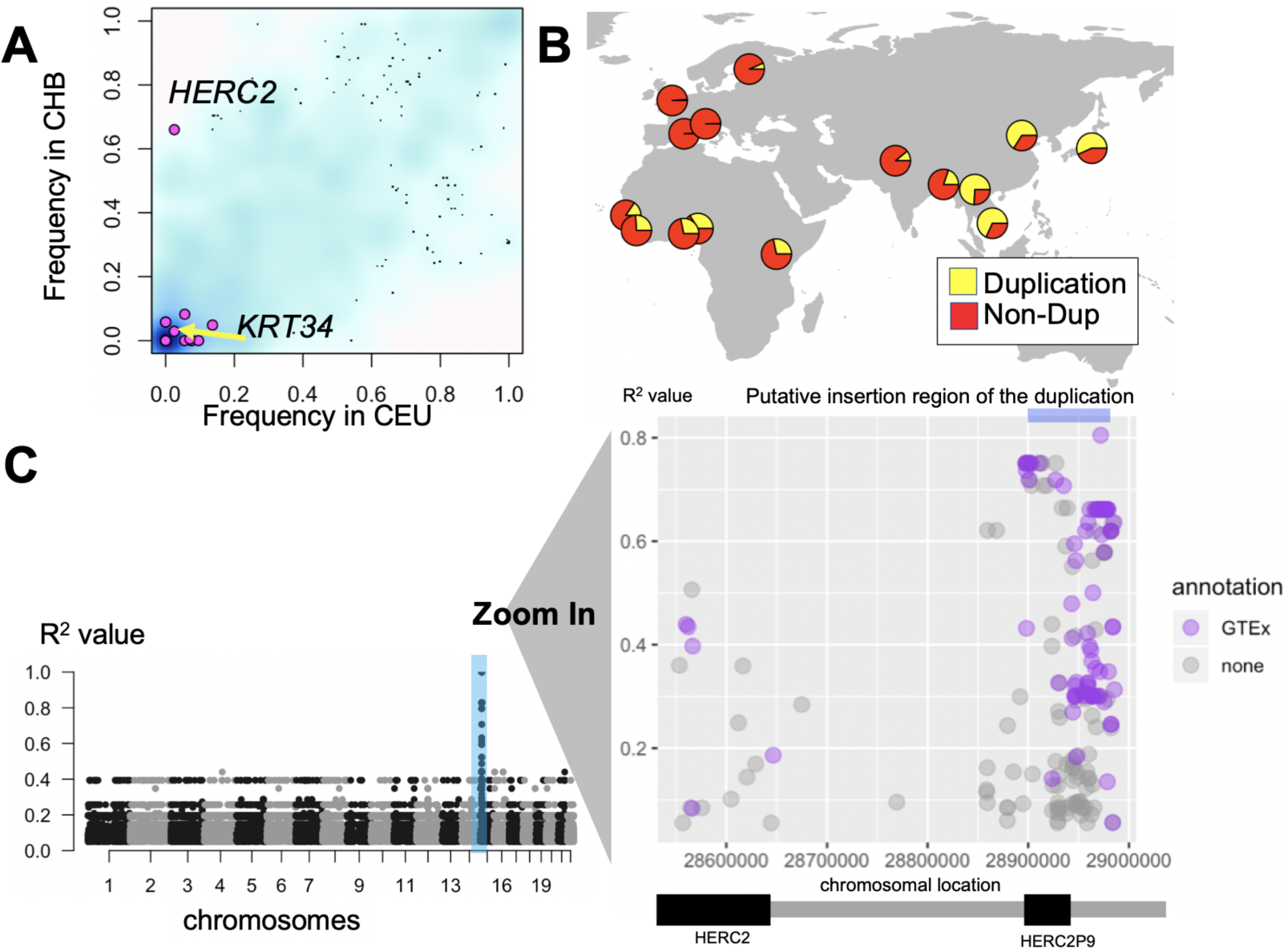
The population differentiation of the partial *HERC2* duplication. **(A)** The frequency of the target duplications which was observed either CHB or CEU (pink dots) and randomly selected 3000 SNPs (> 5% in CEU, YRI or CHB) (blue background cloud). The density of the blue color reflects the density observations. The x-axis shows the frequency of the variation in CEU and the y-axis shows the frequency of the variation in CHB. **(B)** The geographical distribution of the *HERC2* gene duplication allele. Yellow refers to the frequency of duplication allele and red refers to the frequency of the non-duplication allele. **(C)** Left: The putative location of the *HERC2* duplication based on the linkage disequilibrium and the location in the European populations. Right: the magnified version of the left image, chromosome 15. Dots are single nucleotide variants with R^2^ >0.05 with the duplication in this location. The X-axis shows the chromosomal location and Y-axis shows the R^2^ value between the single nucleotide variant and the *HERC2* duplication. The pale blue bar at upper-right indicates the region contains SNPs with high linkage disequilibrium (R^2^ > 0.7) with the *HERC2* duplication, where we assume the insertion point of the duplication (chr15:28898099-28902929). We used this region for the subsequent analysis. The blue colored dots indicate single nucleotide variants that show significant association (p < 0.0001) with expression levels of neighboring genes based on GTEx portal (Lonsdale et al. 2013).

To further characterize this polymorphic duplication, we extended our linkage disequilibrium analysis to include the additional 1000 Genomes populations, categorized across continental meta-populations (see **Materials and Methods**). Based on this analysis, we narrowed down the insertion site of the *HERC2* partial gene duplication to chr15:28898099-28902929, and observed a detectable increase in linkage disequilibrium between the duplication and flanking single nucleotide variants in all three continental populations (**fig. 2C, fig. S2**). We found that linkage disequilibrium was strongest in European populations and weaker in Asian and African populations. To understand the ancestral state of the haplotypes harboring the duplication, we we investigated 17 variants that have strong linkage disequilibrium defining this haplotype (R^2^>0.75 with the duplication in European populations) in chimpanzee, Neanderthal, and Denisovan genomes. This analysis suggests that the duplication is likely derived in the human lineage as compared to chimpanzees. Intriguingly we found that the Neanderthal genome is heterozygous at this locus, carrying both duplicated and non-duplicated haplotypes **(Table S2)**.

Next, we used VCFtoTree (Xu et al. 2017) to obtain an alignment file for the *HERC2* duplication haplotype (hg19: chr15:28898099-28902929), containing 2,504 samples available in the 1000 Genome Phase 3 dataset (Sudmant et al. 2015), as well as the reference chimpanzee genome (The Chimpanzee Sequencing Consortium 2005). We then constructed haplotype networks using PopART (Leigh & Bryant 2015) using the Median Joining Method (Bandelt et al. 1999) (**fig. 3A**). This network reveals an apparent reduction of haplotypic diversity in European populations as compared to East Asian and European populations (**fig. 3B**). This is consistent with the dramatically lower allele frequency of the duplication in European populations, which initially led us to focus on this locus (**fig. 2AB**).

**FIG. 3.**
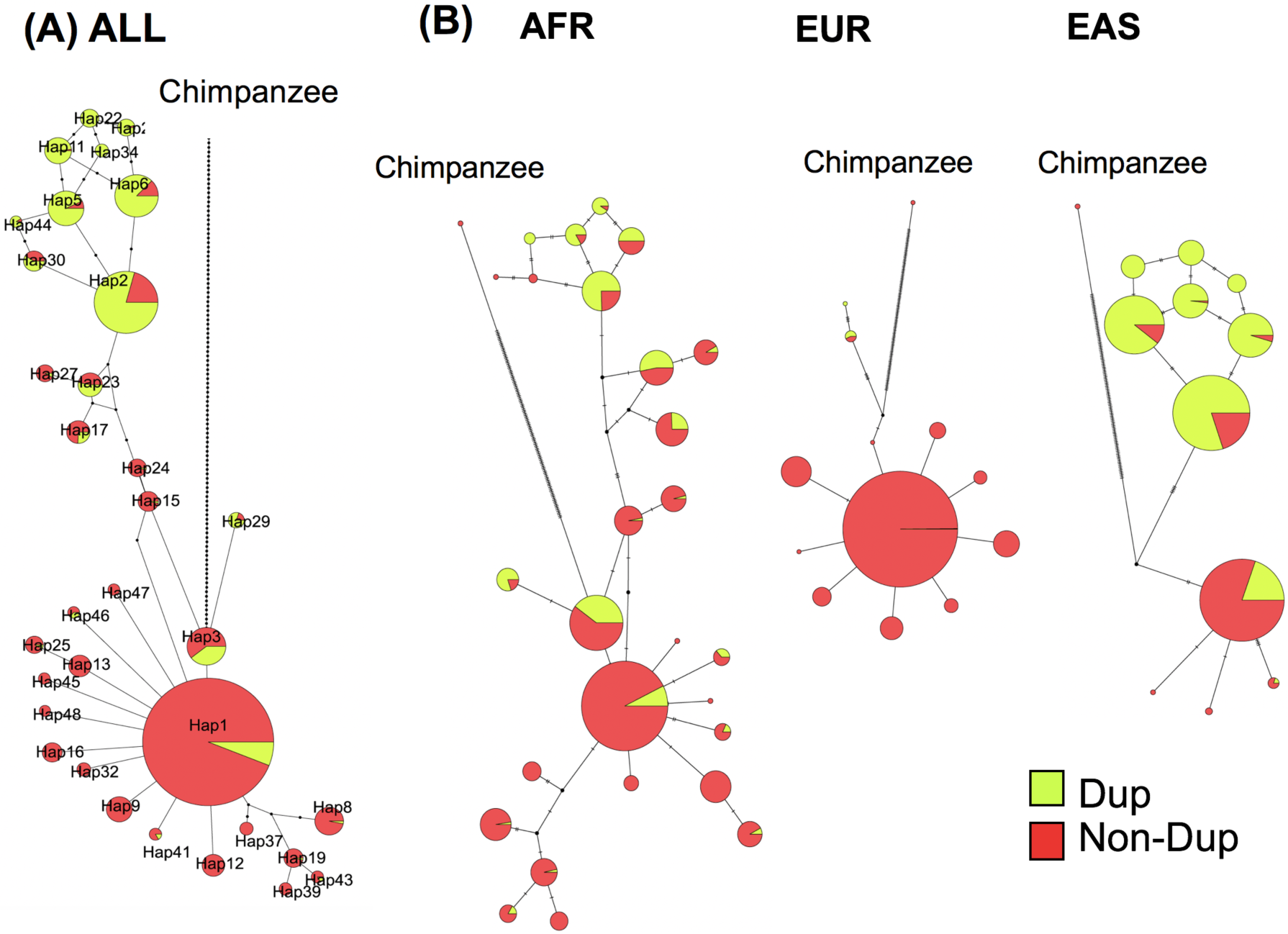
Haplotype networks of the *HERC2* duplication insertion region. **(A)** Merged haplotype network of the three meta-populations (AFR, EUR, EAS) constructed from 3336 haplotypes from chr15:28898099-28902929 (represented in **fig. 2C**). **(B)** Breakdown of individual networks to help visualization of the distribution of alleles in each meta-population. Yellow refers to the frequency of duplication allele and red refers to the frequency of the non-duplication allele.

We then asked whether population-specific selective forces can explain the reduction in haplotypic diversity at this locus in European populations. We calculated several neutrality measures at the locus harboring the *HERC2* duplication and compared these to empirical distributions constructed from 26,283 available 3kb regions across chromosome 15 from the 1000 Genomes Selection Browser (Pybus et al. 2014). We found Tajima’s D scores in this region to be lower than 95% of the observed values on chromosome 15 for European populations (**fig. 4A**). However, in Asian and African populations, Tajima’s D scores fell within the expected range based on the empirical distribution. Tajima’s D measures deviations in the allele frequency spectrum (Tajima 1993); negative values indicate an excess of rare alleles, which may be a consequence of negative or positive selection. In this case, based on network analysis (**fig. 3**), we argue that this signal is primarily driven by the reduction of the allele frequency haplotypes harboring the duplication in the European populations.

**FIG. 4.**
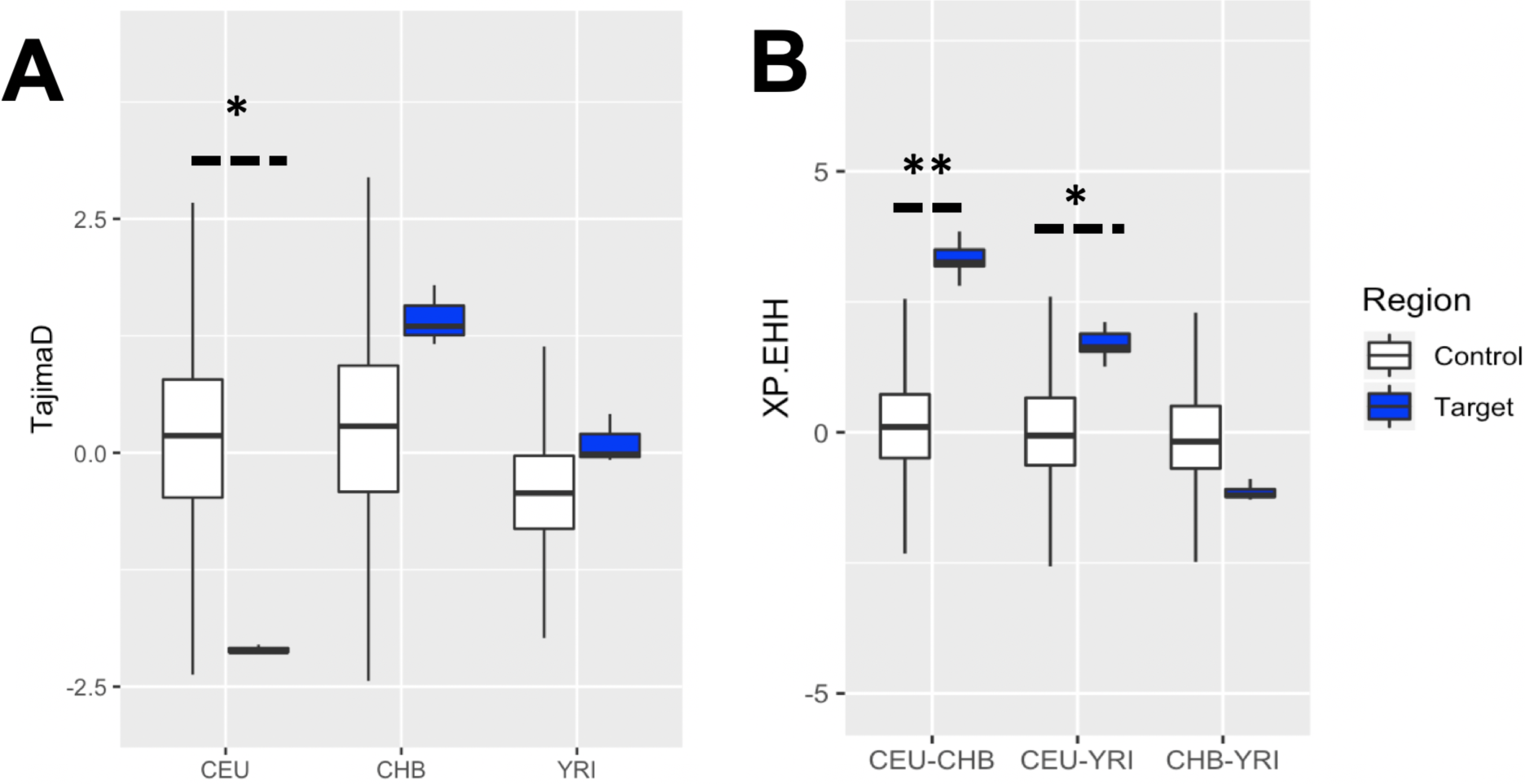
Neutrality test on the putative insertion region of the partial *HERC2* duplication. All values were obtained through the 1000 Genomes selection browser (Pybus et al. 2014). (A) Tajima’s D and (B) XP-EHH values calculated for the *HERC2* target region (chr15:28898099-28902929, represented in **fig. 2C**), compared to the distributions calculated for all the accessible regions on the chromosome 15 on the 1000 Genomes selection browser (Pybus et al. 2014). ** represents more than 99 percentile of the control region; * represents less than 5 or more than 95 percentile of the control region.

Then, we calculated XP-EHH scores between the three representative populations in a pairwise fashion, which we compared to the empirical distribution of XP-EHH values constructed from the same 26,283 randomly chosen regions across chromosome 15 (**fig. 4B**). XP-EHH calculates the probability of runs of homozygosity around a given locus given that there is the same allele between two populations. A significant positive XP-EHH score is indicative of positive selection in the first population, while a negative score indicates positive selection in the second population (Sabeti et al. 2007). Based on this calculation, we found no clear population differentiation between the representative African (YRI) and East Asian (CHB) populations. However, we found the representative European population (Central European from Utah, CEU) to be significantly differentiated from CHB and YRI (>99^th^, and >95^th^ percentile, respectively) (**fig. 4B**). This means that there are relatively long runs of homozygosity in this region in CEU population as compared to CHB and YRI, concordant with the excess of rare variants suggested by Tajima’s D comparisons. In sum, these results are in line with a scenario that a recent selection event in Europe favors non-duplicated haplotypes over duplicated-haplotypes.

Next, we investigated the potential functional impact of the duplication haplotype. We noted that *HERC2* duplication is likely inserted within the neighboring *HERC2P9* gene (**fig. 2C**). It is intriguing that a much more recent duplication of the *HERC2* gene is inserted into an older paralog of the *HERC2*, which is expressed in multiple tissues. It is possible that recombination-based mechanisms facilitated by sequence homology between these genes led to the insertion of the duplication into *HERC2P9*. Eight *HERC2* pseudogenes are reported in Ensembl (Zerbino et al. 2018) distributed across chromosomes 15 and 16, suggesting frequent duplication of this gene. Based on the GTEx portal (Lonsdale et al. 2013), *HERC2P2, HERC2P3*, and *HERC2P9* are expressed, as well as the intact *HERC2* (**fig. S3**).

Furthermore, we found that the duplication haplotype (imputed by rs77868920, R^2^ = 0.75 in European populations) downregulates the expression of *HERC2P9* in various tissues (**fig. 5**). The most significant effect was observed for downregulation was in the sun-exposed skin (*p*-value = 3.3e-17, Normalized effect size= -0.96). It is possible that the polymorphic duplication may affect not only the expression levels but also the transcribed RNA sequence of the *HERC2P2*. This remains an interesting area for further study. While there are multiple SNVs associated with skin color (Crawford et al. 2017) and iris color (Eiberg et al. 2008; Kayser et al. 2008; Sturm et al. 2008) in this region of the genome, the *HERC2* duplication haplotype does not harbor any of them (MacArthur et al. 2017) (**fig. 2C**).

**FIG. 5.**
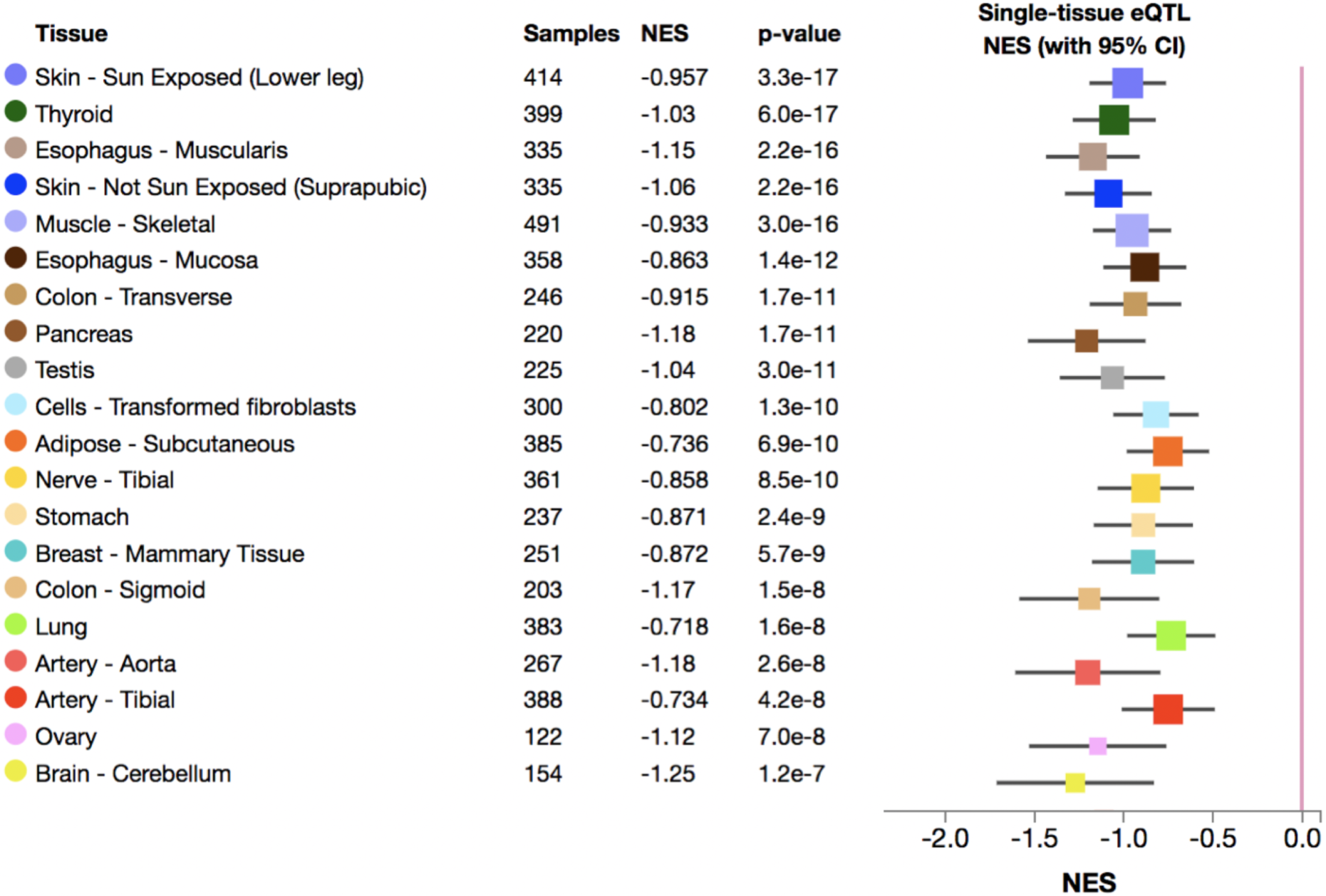
The expression change of the *HERC2P9* gene in various tissues associated with the *HERC2* duplication tag SNP (rs7768920) The top 20 tissues on the GTEx (Lonsdale et al. 2013) with the lowest p-value are shown. Normalized effect size is defined as the slope of the linear regression of the expression of the *HERC2P9* gene for three genotypes of the tag SNP. Normalized effect size is computed as the effect of the alternative allele relative (tagged to duplication) to the reference allele (tagged to non-duplication) in the human genome reference GRCh37/hg19. The whiskers on the plot represent the 95%confidence intervals.

## DISCUSSION

One of the major questions in the population genetics is the impact of polymorphic duplications to phenotypic variation and the evolutionary consequences of this impact. One of the challenges in the field is investigating the associations of the polymorphic duplications to phenotypic variation. Another difficulty is to be able to use standard population genetics tools, which are often designed for analysis of single nucleotide variation. To resolve these problems, we first identified 22 common polymorphic duplications that show linkage disequilibrium with single nucleotide variations across the human genome (**Table 1**). This methodology enabled us to resolve the haplotypes that harbor these duplications. By investigating the single nucleotide variants, we were able to associate two of these haplotypes to skin color and monocyte percentage. Given that these duplications are major mutation events potentially affecting thousands of base pairs, it is likely that they are the causal variants that affect these phenotypes.

Similarly, using single nucleotide variation information, we were able to investigate the evolutionary trajectories of the haplotypes harboring the duplications. Based on such analysis, we here present multiple lines of evidence that the haplotype harboring the partial *HERC2* gene duplication is selected against in European populations. We found that this haplotype affects the expression level of *HERC2P9* significantly, even though the exact phenotype that is under selection is not clear. We argue that as more complete variation datasets and accompanying databases with expression and phenotype data become more available, the haplotype level analysis of gene duplications, in particular, and structural variation, in general, will become more commonplace. Our study represents a first step in integrating multiple data types to understand the evolutionary impact of gene duplications.

## Materials and Methods

### The linkage disequilibrium based detection of duplications

We modified VCFtools (Danecek et al. 2011) to calculate the R^2^ between a target duplication and other variants in a genome-wide manner. We first made a custom genome-wide VCF file from 1000 Genomes phase 3 dataset for CEU, YRI and CHB population. We conducted population-specific analyses to increase the sensitivity of linkage disequilibrium. To reduce file size, we omitted variants which were not observed in the population of interest. Then we calculated the R^2^ between a target duplication and other variants in a genome-wide manner with VCFtools. We visualized linkage disequilibrium linkage disequilibrium by using R qqman package (**fig. 1B**). To ensure the accuracy of these haplotypes, we manually verified the informative variants in the Integrated Genome Browser (Thorvaldsdóttir et al. 2013). For example, we verified one insertion-deletion polymorphism that is in strong linkage disequilibrium (R^2^ = 0.75) with a duplication (esv3635993), which provides a clear example of a likely true-positive variant calling in this region tagging the duplication polymorphism (**fig. S4**).

### The detection of genic/exonic duplications

We used NCBI RefSeq track on UCSC Genome Table Browser to get the gene and exon information. By using Bedtools intersect (Quinlan & Hall 2010), we counted the number of duplications that overlap with i) entire genes, ii) entire exons, iii) more than one base pair of a gene (including introns). Note that none of the 22 duplications that we scrutinized here partially overlap with a coding exon, i.e., if a duplication overlap with a coding sequence, it contains at least one entire exon (**Table 1**). The gene functions listed in Table S1 are based on the genetic associations in Gene Atlas UK Biobank (Canela-Xandri et al. 2018).

We found one transchromosomal duplication, esv3631000, which contains *ZNF664*. To verify this, we carefully checked all the 19 single nucleotide variants that have strong linkage disequilibrium (> 0.8) with this duplication. We found that 17 of those cluster in the 50 kb region on chr2 (chr2:3918719-3970271). The other 2 single nucleotide variants actually overlap with the original duplicated copy on chromosome 12. We thought that these may be false positive calls due to misalignment of the reads originating from the duplicated copy onto the original gene. If this is the case, we expect that the mapped reads to have a ⅓ ratio in a sample where there is a heterozygous duplication. We also expect that these single nucleotide variants are called as heterozygous in all cases. Indeed, we found that 235 out of 2504 individuals are heterozygous for both of these two single nucleotide variants (rs80197353, rs78005948) and no homozygous variants were documented. Furthermore, we manually inspected these single nucleotide variants using exome data from 1,000 Genomes dataset and found that reads carrying the nonreference alleles were found in approximately ⅓ of the reads for both single nucleotide variants. This is not consistent with the expected 50–50 ratio for heterozygous variant calls (**fig. S5**). Collectively, our analysis suggests that the single nucleotide variants on chromosome 2 are likely false positive variant calls due to erroneous read mapping and that the duplication insertion site is indeed on chromosome 12.

### Getting random control regions

To obtain random single nucleotide variants which match our initial filtering process for polymorphic duplications (5%> in CEU, YRI or CHB), we first used bedtools (Quinlan & Hall 2010) for constructing random chromosomal coordinates. We then applied the random chromosomal coordinates to the 1000 Genome variants (Sudmant et al. 2015) and used vcftools (Danecek et al. 2011) to retrieve the allele frequency information. We finally used 3,000 SNPs for the comparison between duplicated regions and random SNPs (**fig. 2A**). In a similar way, we used all the available coordinates on chromosome 15, on which the *HERC2* locates, for the neutrality test on the selection browser (Pybus et al. 2014)

### Population genetics analyses on the *HERC2* duplication

To increase the sensitivity and confirm the initial linkage disequilibrium calculation, we extended the linkage disequilibrium analysis on the original *HERC2* gene - putative *HERC2* duplication region (chr15:28549579-28985567) from CEU to European populations (Utah residents with Northern and Western European ancestry (CEU), Toscani in Italy (TSI), Finnish in Finland (FIN), British in England and Scotland (GBR)), Iberian populations in Spain (IBS)), YRI to African populations (Gambian in Western Division, The Gambia (GWD), Mende in Sierra Leone (MSL), Esan in Nigeria (ESN), Yoruba in Ibadan, Nigeria (YRI), Luhya in Webuye, Kenya (LWK)), CHB to East Asian populations (Han Chinese in Bejing, China (CHB), Japanese in Tokyo, Japan (JPT), Southern Han Chinese, China (CHS), Chinese Dai in Xishuangbanna, China(CDX), Kinh in Ho Chi Minh City, Vietnam (KHV)). We observed similar peaks in all populations in chr15:28898099-28902929 (**fig. 2C, fig. S6**)

### Neutrality Tests

Tajima’s D (Tajima 1993) and XP-EHH (Sabeti et al. 2007) values were calculated by the 1000 Genomes selection browser (Pybus et al. 2014) for the bins contain the target region (chr15:28898099-28902929) and control region (all the available 26,283 available 3kb regions across the chromosome 15).

### Haplotype network analysis

To draw the haplotype networks, we first converted the target region vcf file (chr15:28898099-28902929) from the 1000 Genome phase3 dataset and hg19 reference genome to a fasta file by VCTtoTree (Xu et al. 2017). We also included the chimpanzee genome sequence (The Chimpanzee Sequencing Consortium 2005). We manually checked informative alleles in Neanderthal and Denisovan genomes (Prüfer et al. 2014; Reich et al. 2010). We used Popart (Leigh & Bryant 2015) for the visualization.

**Table 1.** All the 22 duplications and one of their tag SNPs, R^2^ value, and the phenotypic information by Gene ATLAS.

## Supporting information

## Data Availability

Supplementary figures and tables are available online.

## Acknowledgments

This study is supported by OG’s funds from National Science Foundation Grant # 1714867. MS is funded by Astellas Foundation for Research on Metabolic Disorders. We would like to thank Izzy Starr, Dr. Rebecca Torene Iskow and Dr. Yoko Satta for careful reading of this manuscript.

## Supplementary Material

**Figure S1.**
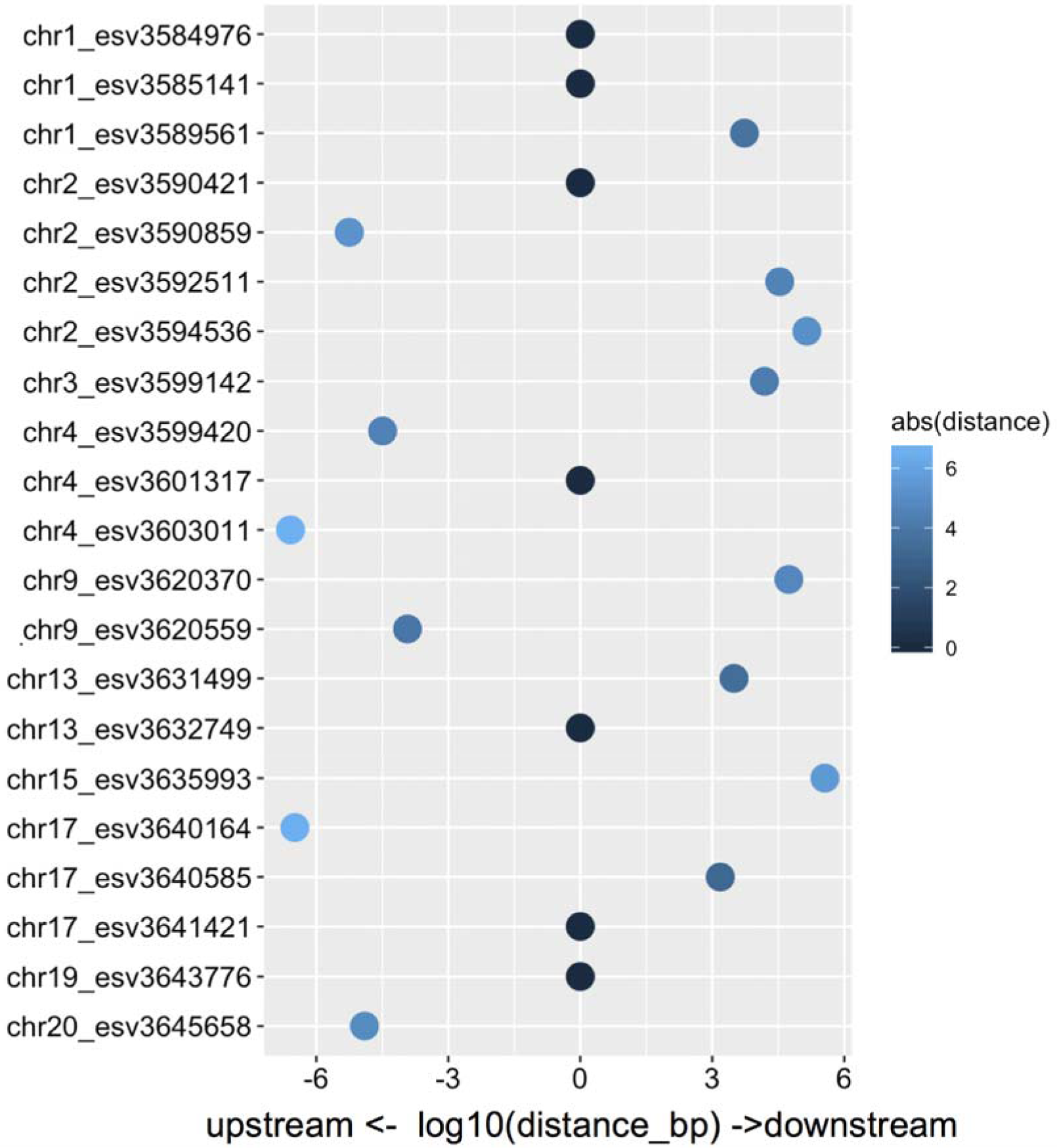
The distance of tag SNP of the duplicates from the original sequence (data from **table 1**). Y-axis is each duplication except for one trans-chromosomal duplicate. X-axis indicates the log_10_ distance (bp) of the tag SNP (the putative insertion region of the newly duplicate) from the original gene. Left is upstream and right is downstream to the original gene. Color corresponds with the distance.

**Figure S2.**
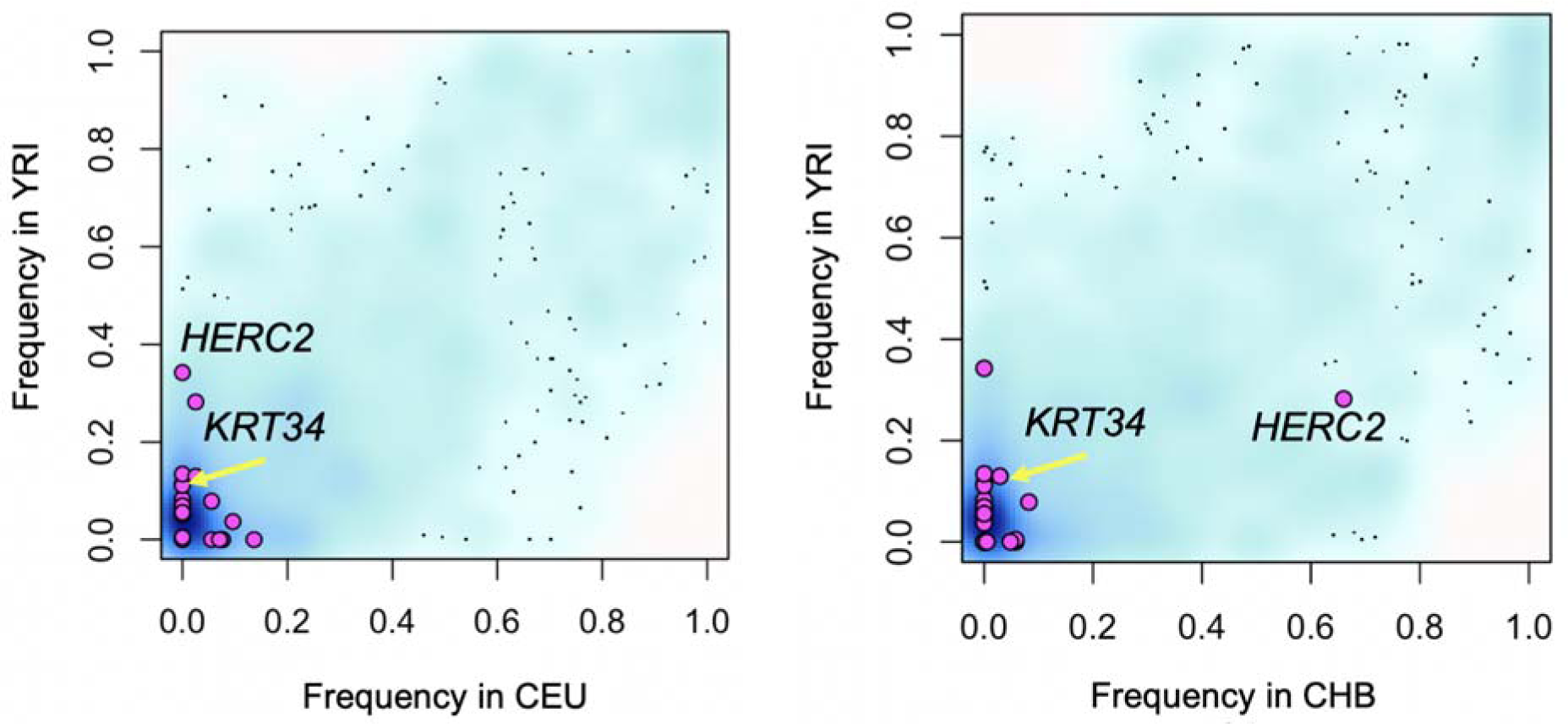
The population allele frequencies of the polymorphic duplications. The allele frequencies of the 22 duplications investigated in this study are shown as pink dots over a background of the allele frequencies of randomly selected 3000 single nucleotide variants (> 5% in CEU, YRI or CHB) (blue cloud). The density of blue color reflects the number of single nucleotide variants.

**Figure S3.**
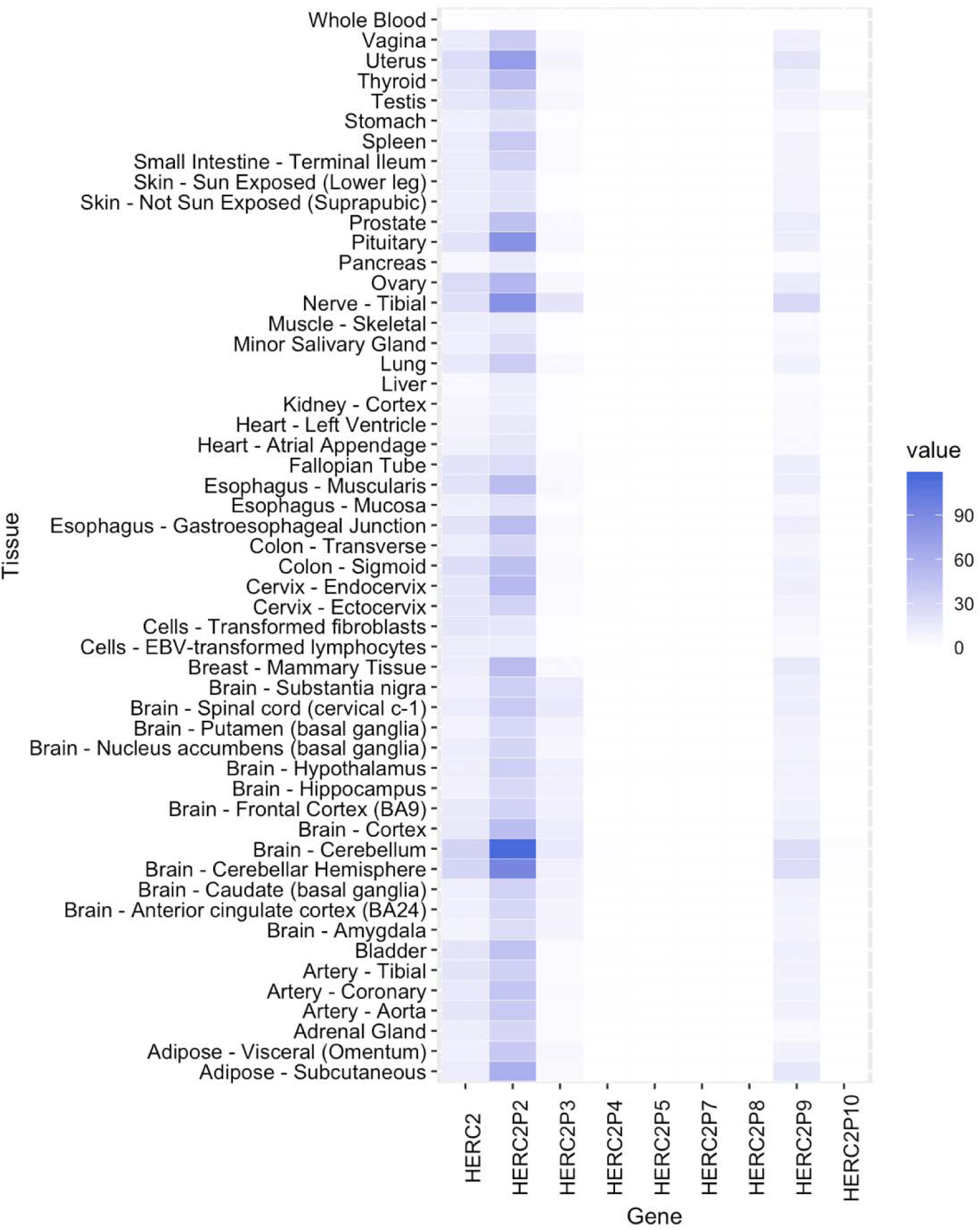
The normalized expression of *HERC2* and its pseudogenes in various tissues (TPM, Transcripts per Million (TPM)) (Li et al. 2010)). Data were obtained at GTEx portal (Lonsdale et al. 2013).

**Figure S4.**
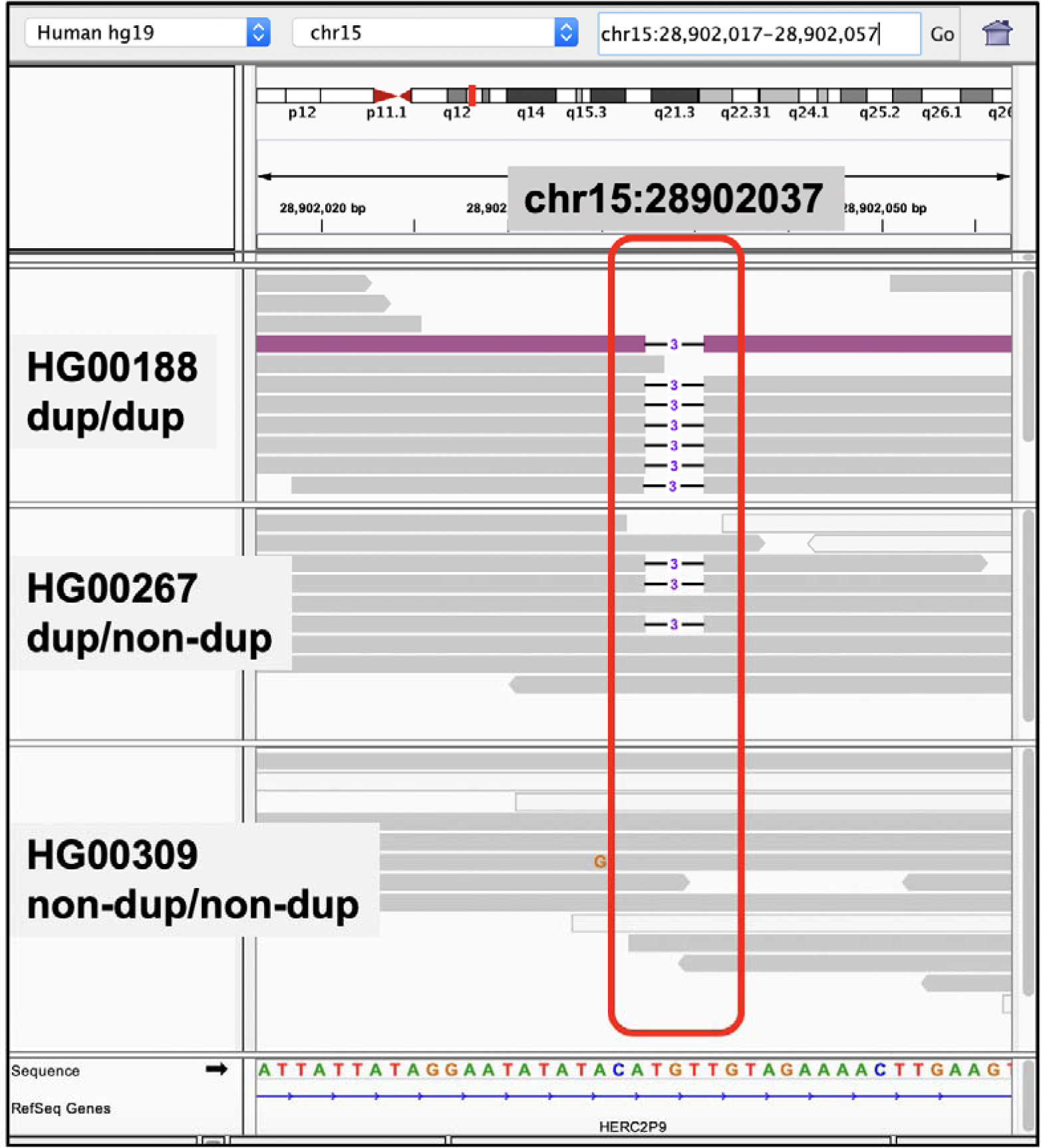
One of the tag variants with the *HERC2* duplication (esv3635993), an insertion-deletion polymorphism (rs202166422) that is in strong linkage disequilibrium (R^2^ = 0.75) with the *HERC2* duplication. The deletion allele of rs202166422 is in the linkage disequilibrium with the duplication allele of esv3635993.

**Figure S5.**
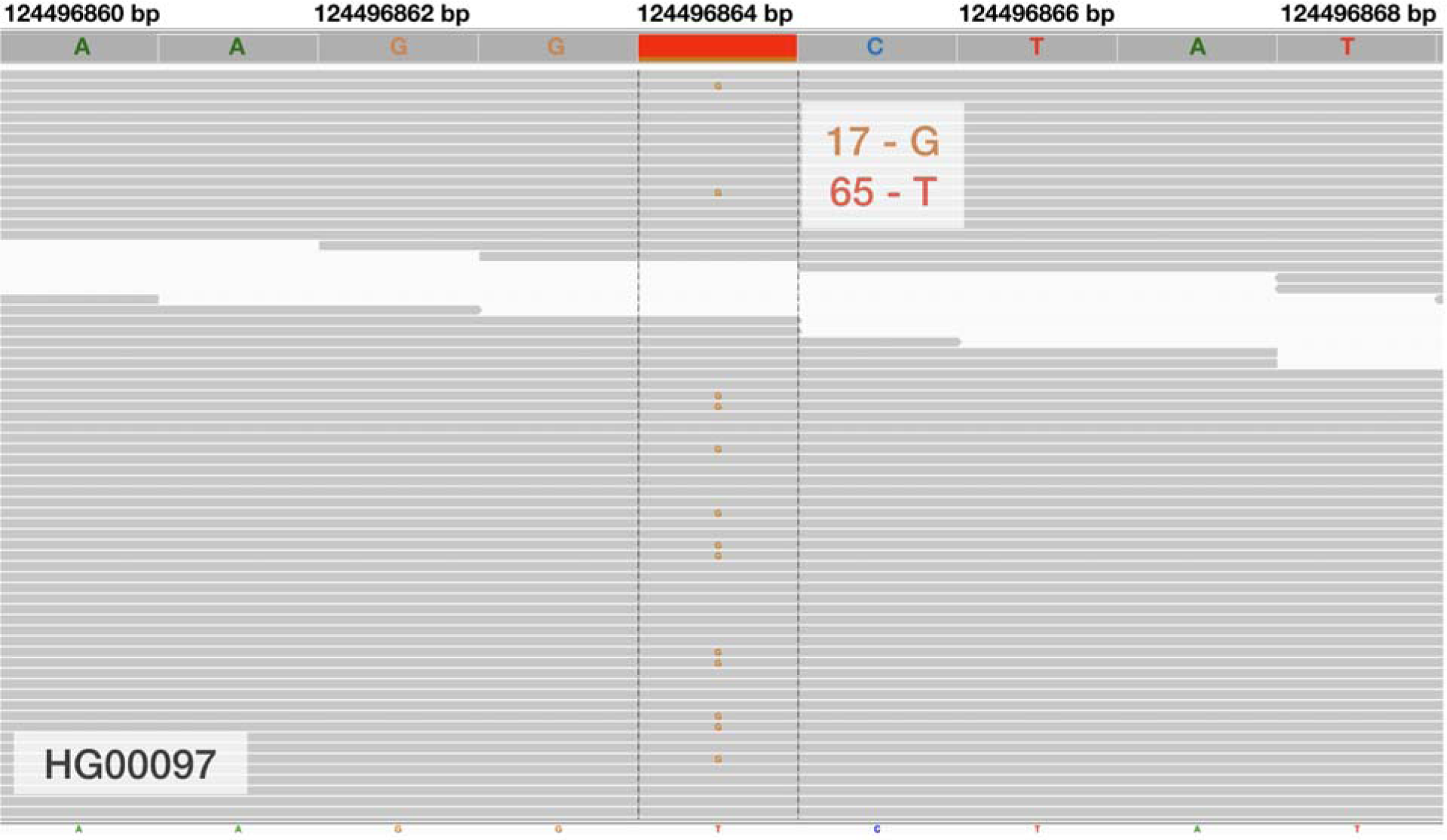
The putatively false SNP call of a candidate tag SNP (rs80197353) of esv3631000 on chromosome 12. We manually inspected these single nucleotide variants using exome data from 1,000 Genomes dataset and found that reads carrying the non-reference alleles were found in 17/82 of the reads for both single nucleotide variants. Our analysis suggests that the single nucleotide variants on chromosome 2 are likely false positive variant calls due to erroneous read mapping and that the duplication insertion site is indeed on chromosome 12.

**Figure S6.**
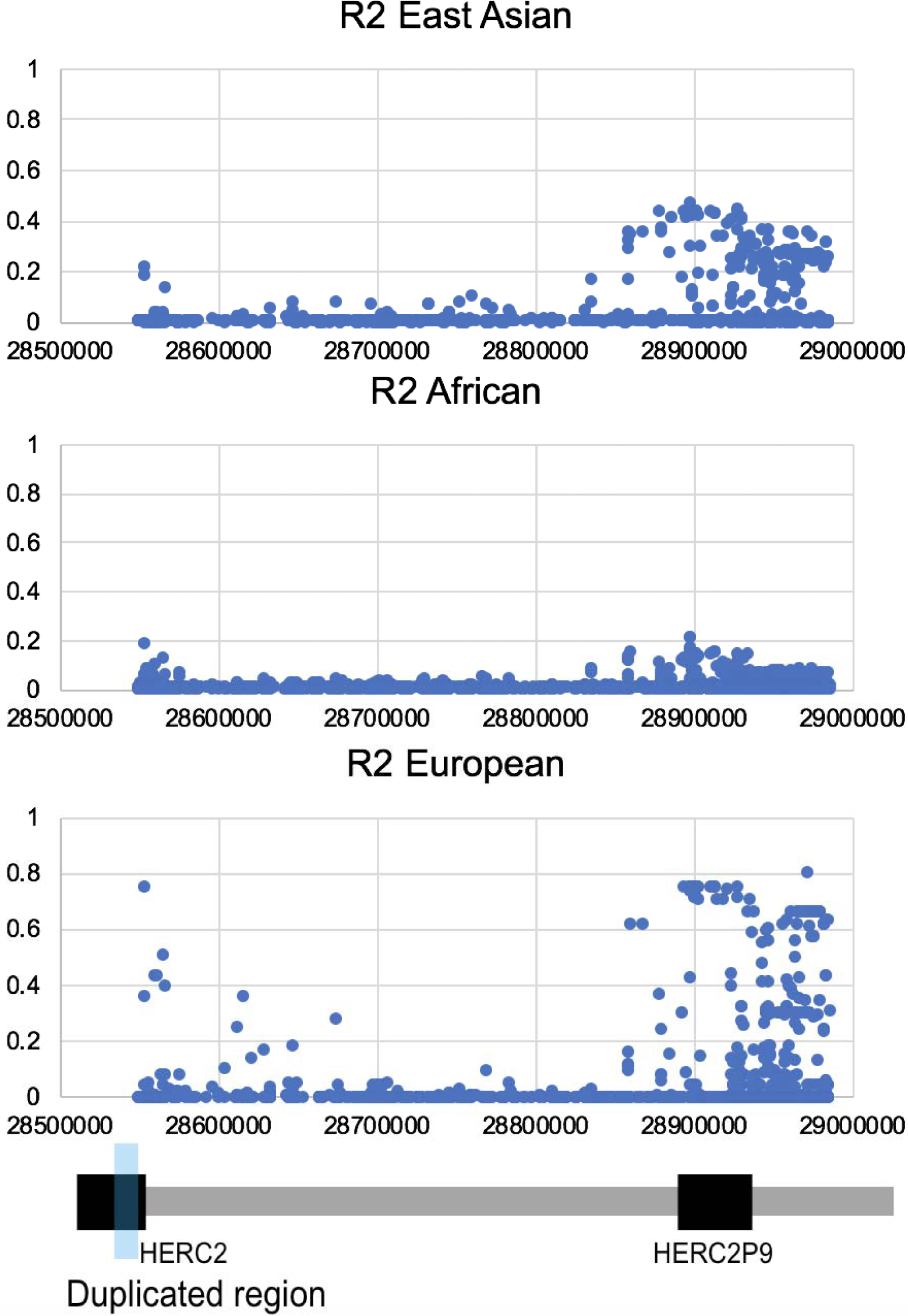
The linkage disequilibrium between the *HERC2* duplication and single nucleotide variants in multiple populations. X-axis: coordinate on chromosome 17, Y-axis: R^2^ between the HERC2 duplication and single nucleotide variants in the three meta-populations in the 1000 Genomes phase 3 dataset.

**Table S1.** Tag SNPs, linkage disequilibrium, observed populations on the investigated 22 polymorphic duplications.

**Table S2.** Alleles of the *HERC2* partial duplication tagging SNPs in Neanderthal, Denisova and reference and alternative alleles in the modern human populations.

## References

Bandelt HJ, Forster P, Röhl A. 1999. Median-joining networks for inferring intraspecific phylogenies. Mol. Biol. Evol. 16:37–48.

Canela-Xandri O, Rawlik K, Tenesa A. 2018. An atlas of genetic associations in UK Biobank. Nat. Genet. 50:1593–1599.

Conrad DF et al. 2010. Origins and functional impact of copy number variation in the human genome. Nature. 464:704–712.

Crawford NG et al. 2017. Loci associated with skin pigmentation identified in African populations. Science. 358. doi: 10.1126/science.aan8433.

Danecek P et al. 2011. The variant call format and VCFtools. Bioinformatics. 27:2156–2158.

Eaaswarkhanth M et al. 2016. Atopic Dermatitis Susceptibility Variants in Filaggrin Hitchhike Hornerin Selective Sweep. Genome Biol. Evol. 8:3240–3255.

Eaaswarkhanth M, Pavlidis P, Gokcumen O. 2014. Geographic distribution and adaptive significance of genomic structural variants: an anthropological genetics perspective. Hum. Biol. 86:260–275.

Eiberg H et al. 2008. Blue eye color in humans may be caused by a perfectly associated founder mutation in a regulatory element located within the HERC2 gene inhibiting OCA2 expression. Hum. Genet. 123:177–187.

Giesen M et al. 2011. Ageing processes influence keratin and KAP expression in human hair follicles. Exp. Dermatol. 20:759–761.

Kayser M et al. 2008. Three genome-wide association studies and a linkage analysis identify HERC2 as a human iris color gene. Am. J. Hum. Genet. 82:411–423.

Leffler EM et al. 2017. Resistance to malaria through structural variation of red blood cell invasion receptors. Science. 356. doi: 10.1126/science.aam6393.

Leigh JW, Bryant D. 2015. popart: full-feature software for haplotype network construction. Methods Ecol. Evol. 6:1110–1116.

Li B, Ruotti V, Stewart RM, Thomson JA, Dewey CN. 2010. RNA-Seq gene expression estimation with read mapping uncertainty. Bioinformatics. 26:493–500.

Lonsdale J et al. 2013. The Genotype-Tissue Expression (GTEx) project. Nat. Genet. 45:580–585.

MacArthur J et al. 2017. The new NHGRI-EBI Catalog of published genome-wide association studies (GWAS Catalog). Nucleic Acids Res. 45:D896–D901.

Meisler MH, Ting CN. 1993. The remarkable evolutionary history of the human amylase genes. Crit. Rev. Oral Biol. Med. 4:503–509.

Mills RE et al. 2011. Mapping copy number variation by population-scale genome sequencing. Nature. 470:59–65.

Pajic P, Lin Y-L, Xu D, Gokcumen O. 2016. The psoriasis-associated deletion of late cornified envelope genes LCE3B and LCE3C has been maintained under balancing selection since Human Denisovan divergence. BMC Evol. Biol. 16:265.

Perry GH et al. 2007. Diet and the evolution of human amylase gene copy number variation. Nat. Genet. 39:1256–1260.

Prüfer K et al. 2014. The complete genome sequence of a Neanderthal from the Altai Mountains. Nature. 505:43–49.

Pybus M et al. 2014. 1000 Genomes Selection Browser 1.0: a genome browser dedicated to signatures of natural selection in modern humans. Nucleic Acids Res. 42:D903–9.

Quinlan AR, Hall IM. 2010. BEDTools: a flexible suite of utilities for comparing genomic features. Bioinformatics. 26:841–842.

Reich D et al. 2010. Genetic history of an archaic hominin group from Denisova Cave in Siberia. Nature. 468:1053–1060.

Sabeti PC et al. 2007. Genome-wide detection and characterization of positive selection in human populations. Nature. 449:913–918.

Sabeti PC et al. 2005. The case for selection at CCR5-??32. PLoS Biol. 3:1963–1969.

Slentz-Kesler KA, Hale LP, Kaufman RE. 1998. Identification and characterization of K12 (SECTM1), a novel human gene that encodes a Golgi-associated protein with transmembrane and secreted isoforms. Genomics. 47:327–340.

Sturm RA et al. 2008. A single SNP in an evolutionary conserved region within intron 86 of the HERC2 gene determines human blue-brown eye color. Am. J. Hum. Genet. 82:424–431.

Sudmant PH et al. 2015. An integrated map of structural variation in 2,504 human genomes. Nature. 526:75–81.

Tajima F. 1993. Simple methods for testing the molecular evolutionary clock hypothesis. Genetics. 135:599–607.

The 1000 Genomes Project Consortium et al. 2015. A global reference for human genetic variation. Nature. 526:68.

The Chimpanzee Sequencing Consortium. 2005. Initial sequence of the chimpanzee genome and comparison with the human genome. Nature. 437:69–87.

Thorvaldsdóttir H, Robinson JT, Mesirov JP. 2013. Integrative Genomics Viewer (IGV): high-performance genomics data visualization and exploration. Brief. Bioinform. 14:178–192.

Weischenfeldt J, Symmons O, Spitz F, Korbel JO. 2013. Phenotypic impact of genomic structural variation: insights from and for human disease. Nat. Rev. Genet. 14:125–138.

Xu D, Jaber Y, Pavlidis P, Gokcumen O. 2017. VCFtoTree: a user-friendly tool to construct locus-specific alignments and phylogenies from thousands of anthropologically relevant genome sequences. BMC Bioinformatics. 18:426.

Zerbino DR et al. 2018. Ensembl 2018. Nucleic Acids Res. 46:D754–D761.

Zhang F, Gu W, Hurles ME, Lupski JR. 2009. Copy number variation in human health, disease, and evolution. Annu. Rev. Genomics Hum. Genet. 10:451–481.

